# Direct binding to RPA-coated ssDNA allows recruitment of the ATR activator TopBP1 to sites of DNA damage

**DOI:** 10.1101/050013

**Authors:** Julyana Acevedo, Shan Yan, Matthew Michael

**Author notes:** To whom correspondence should be addressed: Prof. W. Matthew Michael, Molecular and Computational Biology Section, Department of Biological Sciences, University of Southern California, Room 104B RRI, 1050 Childs Way, Los Angeles, CA 90089.

## Abstract

A critical event for the ability of cells to tolerate DNA damage and replication stress is activation of the ATR kinase. ATR activation is dependent on the BRCT repeat-containing protein TopBP1. Previous work has shown that recruitment of TopBP1 to sites of DNA damage and stalled replication forks is necessary for downstream events in ATR activation, however the mechanism for this recruitment was not known. Here, we use protein binding assays and functional studies in *Xenopus* egg extracts to show that TopBP1 makes a direct interaction, via its BRCT2 domain, with RPA-coated ssDNA. We identify a point mutant that abrogates this interaction, and show that this mutant fails to accumulate at sites of DNA damage, and that the mutant cannot activate ATR. These data thus supply a mechanism for how the critical ATR activator, TopBP1, senses DNA damage and stalled replication forks to initiate assembly of checkpoint signaling complexes.

## EXPERIMENTAL PROCEDURES

### *Xenopus* egg extracts and sperm chromatin isolation

Egg extracts and sperm chromatin were prepared as described (1). Sperm chromatin was isolated from egg extract as described (1).

### Expression vectors and IVT protein production

All of the expression vectors used for IVT protein production were based on pCS2+MT. All cloning was done according to standard procedures. The TopBP1 cDNA used for our experiments was isoform TopBP1-B (NCBI accession number AAP03894), cloned into pCS2+MT (2). This protein was initially named Xmus101 (2). All deletion mutants were produced by PCR, using TopBP1-B cDNA as template. PCR primers included an NcoI site on the 5′ end and an XhoI site on the 3′end. Fragments were subcloned into NcoI/XhoI digested pCS2+MT, and verified by DNA sequencing. For the TopBP1 deletion mutants, the amino acid sequence for each deletion mutant is given below: BRCT0-5 (aa 1- 758); BRCT6-8 (aa 759- 1513); BRCT0-2 (aa 1-333); BRCT3 (aa 333- 480); BRCT4-5 (aa 480-758); BRCT0-1 (aa 1-191); BRCT1 (aa 99- 191); BRCT1-2 (aa 99-333); BRCT2 (aa 191-333); BRCT4-5 (aa 527-800). The TopBP1 Δ4-5 mutant was made by deleting aa 588-710. The W265R point mutant was previously described (1). IVT reactions were performed using a TnT^®^ SP6 Quick Coupled Transcription/Translation System, according to the vendor’s instructions (Promega).

### Recombinant proteins

Expression vectors encoding the *Xenopus* RPA trimer and the Rad9 tail domain were kind gifts of K. Cimprich. *Xenopus* RPA trimer was expressed in *E. coli* BL21 cells and purified under denaturing conditions using urea. Proteins were then renatured using sequential dialysis in buffers containing decreasing concentrations of urea. Proteins were then purified via nickel-agarose chromatography. A detailed protocol for the purification is available upon request. All GST fusion proteins were expressed and purified from *E. coli* using standard conditions.

### ssDNA binding assays

In brief, biotin-linked DNA fragments of varying sizes were produced by PCR and then denatured after coupling to Dynabeads^®^ Streptavidin (Life Technologies). Beads were mixed with egg extract or IVT proteins and binding buffer, isolated, washed in binding buffer, and eluted with 2x Laemmli Sample Buffer (2XSB). Biotin-dsDNA was coupled to magnetic streptavidin beads in BW buffer (5mM Tris, pH7.4, 1M NaCl and 0.5mM EDTA), and optionally treated with 0.15N NaOH to produce ssDNA. Binding reactions contained either egg extract, IVT proteins, or purified proteins mixed optionally with purified RPA and dsDNA or ssDNA beads in Buffer A (10mM Hepes, pH7.6, 80mM NaCl, 20mM B-glycerol phosphate, 2.5mM EGTA and 0.1% NP-40). Reaction volumes were 60 ul. Reactions were incubated for an hour at room temperature, the beads were isolated on a magnetic stand, and washed three times with 500 ul of Buffer A. Beads were then eluted in 2X Laemmle sample buffer.

### Immunodepletion and antibodies

RPA and TopBP1 were depleted from *Xenopus* egg extract as described (1). TopBP1 antibodies have been described (2). RPA antibodies were a kind gift of J. Walter. Monoclonal antibody 9E10 (Sigma) was used to detect myc-tagged IVT proteins, phosphorylated Chk1 was detected with Phospho-Chk1 (Ser345) Antibody #2341 (Cell Signaling Technology), and GST fusion proteins were detected with anti-GST antibodies (Sigma).

## INTRODUCTION

The maintenance of genome stability relies on faithful DNA replication and the ability of cells to suppress the mutagenic consequences of replication stress and DNA damage. Two protein kinases, *<u>a</u>taxia <u>t</u>elangiectasia* <u>m</u>utated (ATM) and <u>AT</u>M and <u>R</u>ad3-related (ATR), are perched atop signaling cascades that control cell cycle progression, DNA repair, replication fork stability, and transcriptional responses to DNA damage and replication stress (reviewed in 3-5). ATM is primarily activated by DNA double-strand breaks (DSBs), whereas ATR is activated by stalled replication forks and DSBs. Upon activation, ATR phosphorylates and activates numerous substrates, including the Chk1 kinase (6,7). Here, we address the mechanism for ATR activation, with a focus on how the critical ATR activator, TopBP1, is recruited to sites of DNA damage.

TopBP1 is a scaffold protein that contains nine copies of the <u>BR</u>CA1 <u>C</u>-<u>t</u>erminus (BRCT) domain. BRCT domains mediate protein-protein and protein-DNA interactions and are heavily represented amongst DNA damage response proteins (reviewed in 8). TopBP1 performs multiple functions in chromosome metabolism (reviewed in 9), including the initiation of DNA replication (2,10) and ATR activation (11). Previous work has shown that the roles of TopBP1 in replication initiation and ATR signaling are distinct (11,12). Unlike simpler eukaryotes such as the budding yeast, where multiple factors including the TopBP1 ortholog Dpb11 can activate ATR (13), in metazoans TopBP1 is the sole ATR activator that has been identified to date.

Previous work has detailed important aspects of how ATR is activated by stalled forks. For this, ATR associates with a binding partner, <u>ATR</u>-<u>i</u>nteracting <u>p</u>rotein (ATRIP), and the complex is localized to stalled forks via a direct interaction between ATRIP and <u>r</u>eplication <u>p</u>rotein <u>A</u> (RPA)-coated <u>s</u>ingle-<u>s</u>tranded DNA (ssDNA) (14-19). RPA-coated ssDNA (RPA-ssDNA) is generated at stalled forks due to the uncoupling of DNA helicase and polymerase activities that occurs when the polymerase stalls but the helicase does not (20). ATRIP-mediated ATR docking is necessary but not sufficient for kinase activation. Independent of ATR-ATRIP, another protein complex forms on RPA-ssDNA, and then joins with ATR-ATRIP to activate ATR (21). This second complex contains, minimally, the Rad9-Rad1-Hus1 (911) trimeric clamp protein, the clamp loader Rad17-<u>r</u>eplication <u>f</u>actor <u>C</u> (RFC), and TopBP1. Upon assembly of this second complex, TopBP1 is altered so that its ATR activation domain is revealed, and this allows interaction between TopBP1 and ATR-ATRIP in a manner that activates ATR kinase (11,22).

TopBP1 and ATR also play critical roles in the cellular response to DSBs in many higher eukaryotes, including *Drosophila, Xenopus, C. elegans*, and humans (23-28). TopBP1 and ATR are not only required for the DSB response in mitotic cells, but also for efficient completion of meiotic recombination (reviewed in 29). The mechanism for ATR activation at sites of DSBs is not fully understood. Work in *Xenopus* has outlined a pathway whereby DSBs activate ATM, which then phosphorylates TopBP1, and this allows TopBP1 to activate ATR (30). Previous work has also shown that, in human cells containing DSBs, ATR activity towards different substrates requires distinct activation modes. While TopBP1 is required for ATR activity in all cases, Rad17 (and by extension the 911 complex) is more important for Chk1 phosphorylation while the Nbs1 protein is more important for ATR-directed phosphorylation of RPA32 (31). A dual involvement of Rad17 and Nbs1 in ATR activation has also been observed in *Xenopus* (32,33).

Given the central role of TopBP1 in ATR signaling at both stalled forks and DSBs it is important to understand how TopBP1 is recruited to these different DNA structures. At stalled forks, TopBP1 can form a complex with 911-Rad17-RFC by virtue of an interaction between the BRCT 1 and 2 domains within TopBP1 and the C-terminal tail domain of Rad9 (34-37). This interaction requires phosphorylation of Rad9 on S373 (in the *Xenopus* protein), a site which is thought to be constitutively phosphorylated by casein kinase II (38). The TopBP1-Rad9 tail interaction is required for the ability of TopBP1 to contact DNA-bound ATR-ATRIP complexes, and for ATR activation (34,35). The discovery of the TopBP1-Rad9 interaction led to a simple model for TopBP1 recruitment, whereby the 911 clamp is loaded by Rad17-RFC, and TopBP1 is then recruited via interaction with the Rad9 tail. Four recent findings, however, have challenged this view, and suggest that initial TopBP1 recruitment to sites of damage is independent of 911.

First, multiple lines of published evidence point to a direct role for TopBP1 in 911 loading. A study from our laboratory has shown that when replicating chromatin is isolated and transferred to *Xenopus* egg extract lacking TopBP1, 911 fails to load onto stalled replication forks (1). Furthermore, chromatin transfer experiments showed that TopBP1 must be physically present with 911 for 911 to load onto stalled forks (1). In addition, a TopBP1 point mutant (W265R) was identified that cannot accumulate at sites of replication stress, and, in egg extract containing this mutant as the sole source of TopBP1, both Rad17 and 911 fail to associate with stalled replication forks and ATR is not activated (1). This mutant is, however, competent to initiate DNA replication (1). A requirement for TopBP1 in 911 recruitment has also been observed in human cells, as siRNA-mediated depletion of TopBP1 prevents recruitment of 911 to chromatin after hydroxyurea treatment (39) or UV irradiation (40). These data make it clear that TopBP1 arrives at the stalled fork either before or concurrent with Rad17 and 911, but not after.

Second, it has been demonstrated that the TopBP1-Rad9 tail interaction is dispensable for initial recruitment of TopBP1 to stalled forks (36). In these studies, TopBP1 was shown to accumulate on stalled forks even when Rad9 tail phosphorylation was prevented by a S373A mutation. Third, in the context of meiotic DSBs in mice, TopBP1 recruitment is unhindered in *Hus1* conditional knock-out spermatocytes (41). Fourth, for ATR activity towards RPA32 at DSBs in human cells, Rad17 is dispensable but TopBP1 is not (31).

These recent findings call for a revision to the simple model where TopBP1 is recruited to sites of damage by virtue of the interaction with the Rad9 tail. Rather, TopBP1 accumulates at stalled forks and DSBs via a previously unknown mechanism, and once it is there then 911 loading and ATR activation follow. In order to fully understand how ATR is activated during replication stress and DNA damage, it is therefore necessary to understand the mechanism for the initial TopBP1 recruitment to stalled forks and DSBs. As detailed below, we have elucidated this mechanism, and it involves direct interaction between the TopBP1 BRCT2 domain and RPA-ssDNA.

## RESULTS

### TopBP1 binds ssDNA in a length-and RPA-dependent manner

Previous studies have revealed that the platform for ATR signaling is the RPA-ssDNA that becomes exposed upon uncoupling of helicase and polymerase activities during replication stress, or after resection of DSBs (reviewed in 4). Given that RPA-ssDNA is a common feature of both stalled forks and DSBs, and that TopBP1 activates ATR in both contexts, we reasoned that interaction between TopBP1 and RPA-ssDNA is likely to be important for how TopBP1 is recruited to sites of damage. We therefore examined the ability of TopBP1 to bind to ssDNA after incubation in *Xenopus* egg extract. Magnetic streptavidin beads were coupled to either buffer or equal amounts of either dsDNA or ssDNA and then incubated in egg extract (Fig. 1A). Following incubation, the beads were isolated, washed, and protein binding was assessed by Western blot. As shown in Fig. 1A, TopBP1 could bind well to ssDNA beads, but did not efficiently bind either dsDNA beads or beads alone. To better characterize TopBP1’s binding to ssDNA, we next determined size requirements for binding. We produced ssDNAs of different lengths and these were incubated in egg extract, recovered, washed, and assayed for protein binding. As shown in Fig. 1B, efficient binding of TopBP1 was observed for ssDNAs of >250 nt; however, binding was not observed for a 100 nt piece of ssDNA. Interestingly, RPA could bind efficiently to the 100 nt ssDNA. These data reveal a size threshold for efficient TopBP1 binding to the ssDNA, between 100 and 250 nt. This data is intriguing since the DNA length dependency observed might be the result of secondary structure formation of ssDNA, that might either affect how tightly RPA binds to it or allows RPA to undergo a different conformation when on DNA of different lengths, as it has been noted that RPA undergoes a couple of conformational changes once on DNA. Further work will be required to fully explain this observation.

**Figure 1.**
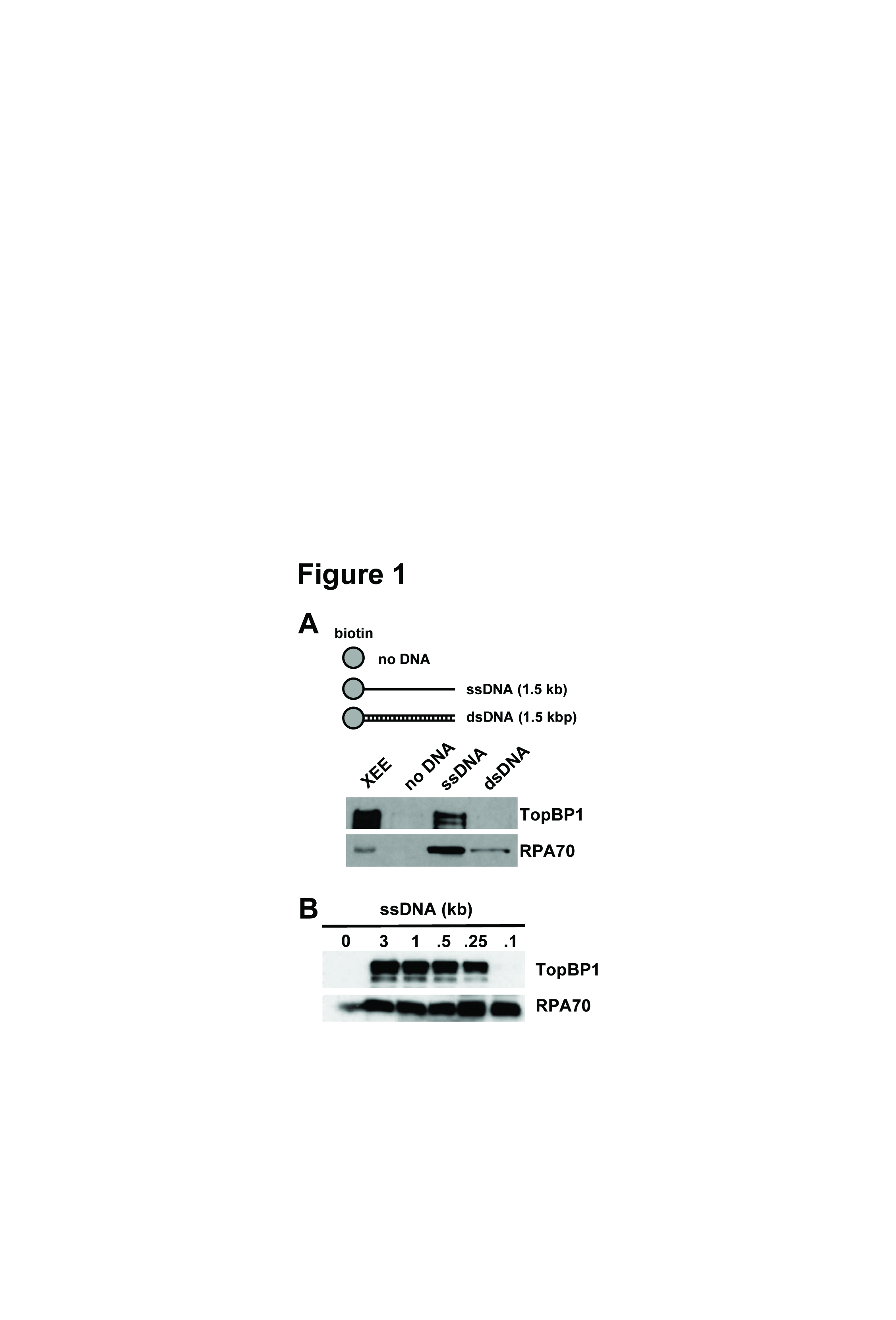
TopBP1 binds ssDNA in a length-dependent manner. (A) The depicted biotin-coupled DNAs were mixed with 50 ul of *Xenopus* egg extract, isolated using streptavidin magnetic beads, washed, eluted, and bound proteins detected by Western blot. Blots were probed with the indicated antibodies. XEE refers to 1 ul of egg extract. (B) Same as (A) except the size of ssDNA was varied, as depicted.

To pursue these observations, we next asked if RPA is required for TopBP1 interaction with ssDNA in egg extract. When RPA was removed from the extract by immunodepletion, TopBP1 could not bind efficiently to ssDNA beads, although binding was observed in the mock-depleted control extract (Fig. 2A). This suggests that RPA is necessary for TopBP1 to bind ssDNA in egg extract. Next, we wanted to determine if RPA is also required for TopBP1 binding to nuclease treated chromatin as seen with ssDNA. We again removed RPA from the extract by immunodepletion and found that RPA-depletion also prevented TopBP1 from binding EcoRI treated chromatin (Fig. 2B). To ask if RPA is sufficient for TopBP1 ssDNA binding, we prepared a recombinant, purified form of the RPA trimer, and used it to prepare RPA-ssDNA templates for TopBP1 binding assays. Recombinant, myc-tagged TopBP1 was produced by in vitro transcription and translation (IVT) in rabbit reticulocyte lysates. IVT TopBP1 was then incubated with purified RPA and ssDNA and assayed for binding. We observed that TopBP1 bound, in an RPA dose-dependent manner, to the ssDNA beads (Fig. 2C). Binding was not observed when RPA was omitted from the reactions. We next asked if TopBP1 could bind to the RPA trimer in the absence of ssDNA. RPA trimer was immobilized on nickel (NTA)-agarose beads by virtue of a 6-histidine tag on RPA70, and incubated with IVT TopBP1. As shown in Fig. 2D, TopBP1 did not bind to immobilized RPA. Based on these data, we conclude that TopBP1 binds to RPA-ssDNA, but not ssDNA alone nor RPA trimer alone. We next asked if there was any difference for TopBP1 binding between RPA-ssDNA formed in egg extract versus RPA-ssDNA formed with purified, recombinant RPA. For this, ssDNA beads were incubated either with egg extract or purified RPA. RPA-ssDNA complexes were then isolated, washed, and incubated with IVT TopBP1. As shown in Fig. 2E, recombinant RPA was just as good as the RPA in egg extract in allowing TopBP1 binding to the ssDNA beads. This shows that any post-translational modifications on RPA that occur in egg extract are dispensable for TopBP1 binding.

**Figure 2.**
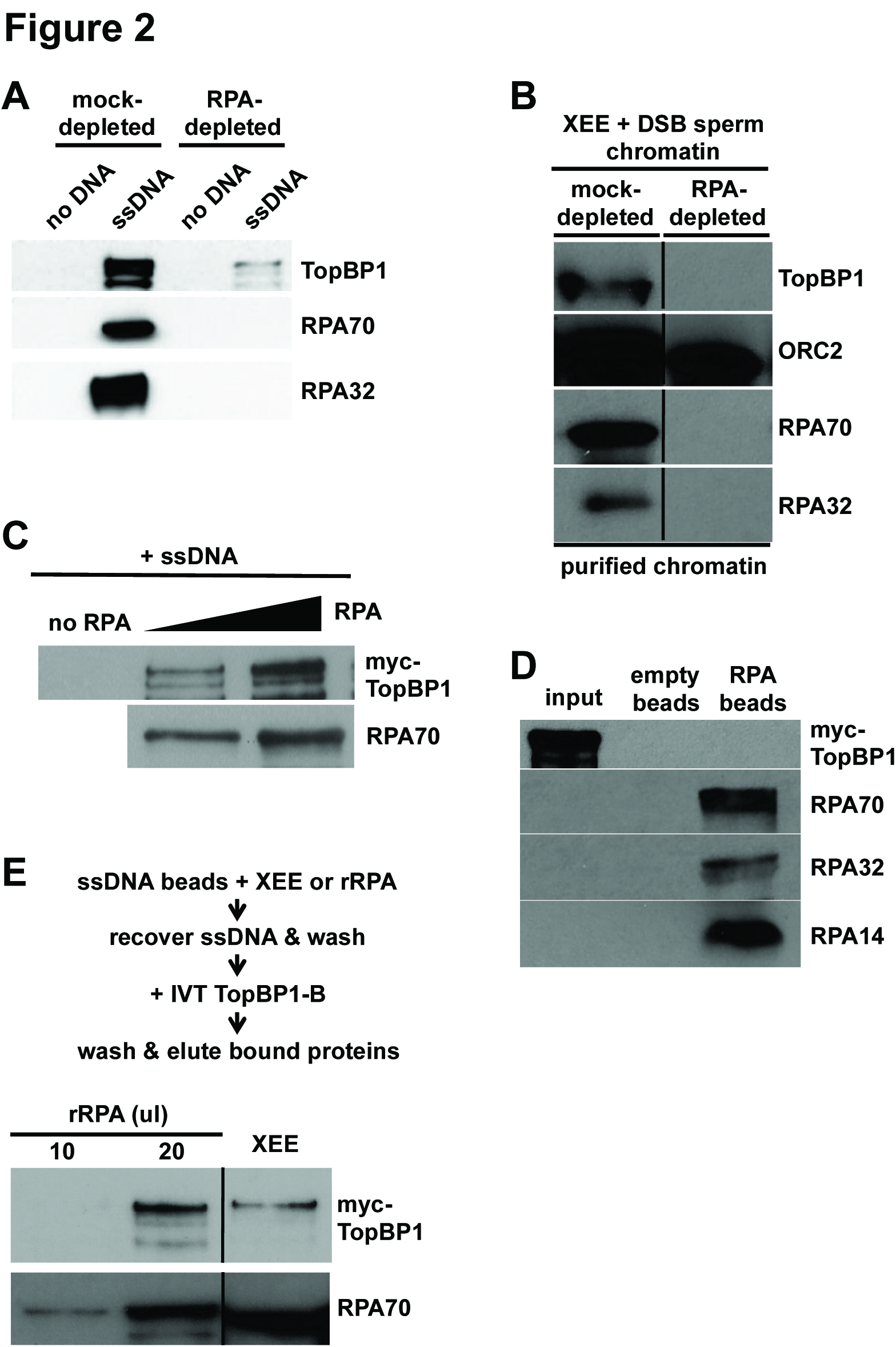
TopBP1 binds RPA-ssDNA, and not ssDNA alone nor free RPA. (A) Same as Fig. 1A except *Xenopus* egg extract was either mock-depleted or depleted of RPA using anti-RPA antibodies, as indicated. (B) *Xenopus* egg extract was either mock-depleted or depleted of RPA using anti-RPA antibodies, as indicated, and TopBP1 binding to EcoRI treated chromatin was assessed. Orc2 serves a loading control and RPA 70kDa and 32kDa shown to confirm immunodepletion. (C) IVT myc-tagged TopBP1 was mixed with varying amounts of purified, recombinant RPA and biotin-coupled ssDNA (1.5 kb). The ssDNAs were isolated using streptavidin magnetic beads, washed, eluted, and bound proteins detected by Western blot. TopBP1 was detected using Mab 9E10, which recognizes the myc tag, and RPA was detected using antibodies against the *Xenopus* RPA trimer. (D) Purified, recombinant RPA trimer was optionally coupled to nickel NTA agarose beads by virtue of a 6-histidine tag on the 70 kDa subunit. Beads were mixed with IVT myc-tagged TopBP1, washed, and bound proteins detected by Western blot. TopBP1 and RPA were detected as in (C). Input refers to 5% of the initial reaction volume. (E) Biotin-linked ssDNA was mixed with either egg extract (XEE) or differeing amounts of purified RPA. The ssDNAs were isolated using streptavidin magnetic beads, washed, and then incubated with IVT myc-tagged TopBP1. The beads were isolated again, washed, and bound proteins detected as in (C). The data shown are from different regions of the same gel, and the images were then spliced together, as indicated by the black line, so as to remove irrelevant material.

### TopBP1 uses its BRCT2 domain to efficiently bind RPA-ssDNA

To further analyze the TopBP1-RPA-ssDNA interaction identified in Figs. 1-2, we used deletion analysis to elucidate the TopBP1 sequence determinants required for binding. TopBP1 is composed of 9 copies of the BRCT domain, termed 0 through 8 (37), that are scattered throughout the length of the protein (Fig. 3A). For our domain delineation experiments, myc-tagged fragments of TopBP1 were produced by IVT and then incubated with ssDNA and, optionally, purified recombinant RPA. Binding was then assessed by Western blot. Bifurcation of the protein into N-terminal and C-terminal portions revealed that the N-terminal half, containing BRCTs 0-5, bound ssDNA in an RPA-dependent manner, whereas the C-terminal half did not (Fig. 3B). Subdivision of the N-terminal half revealed that fragments containing BRCTs 0-2 and 4-5 bound ssDNA in an RPA-dependent manner, whereas a fragment containing BRCT3 did not (Fig. 3C). Further analysis of the region containing BRCTs 0-2 revealed that BRCT2 alone can bind efficiently to ssDNA, in an RPA-dependent manner, whereas fragments containing either BRCTs 0-1 or BRCT1 alone could not (Figs. 3D&E). These data reveal that BRCT2 binds well to RPA-ssDNA. We next further analyzed the region containing BRCT4-5, and here the results were surprising. BRCT4 bound very well to both ssDNA and RPA-ssDNA, and BRCT5 showed a similar pattern, albeit with reduced binding relative to BRCT4 (Fig. 3F). In this experiment, BRCT0-2 bound specifically to RPA-ssDNA, as did BRCT4-5 (Fig. 3F). These data show that when BRCT domains 4 and 5 are contiguous then binding to ssDNA requires RPA, however when the domains are separated from one another then the requirement for RPA is lost. The basis for this is not known. To assess the importance of the BRCT4-5 domains for binding of TopBP1 to RPA-ssDNA, we tested an internal deletion mutant that lacks these sequences (TopBP1 Δ4&5). As shown in Fig. 3G, this protein bound as well as its wild type counterpart to RPA-ssDNA. This experiment shows that when the remainder of the protein is present then BRCT domains 4-5 do not contribute much to binding to RPA-ssDNA, and thus for the remainder of this study attention was focused on the interaction between BRCT2 and RPA-ssDNA.

**Figure 3.**
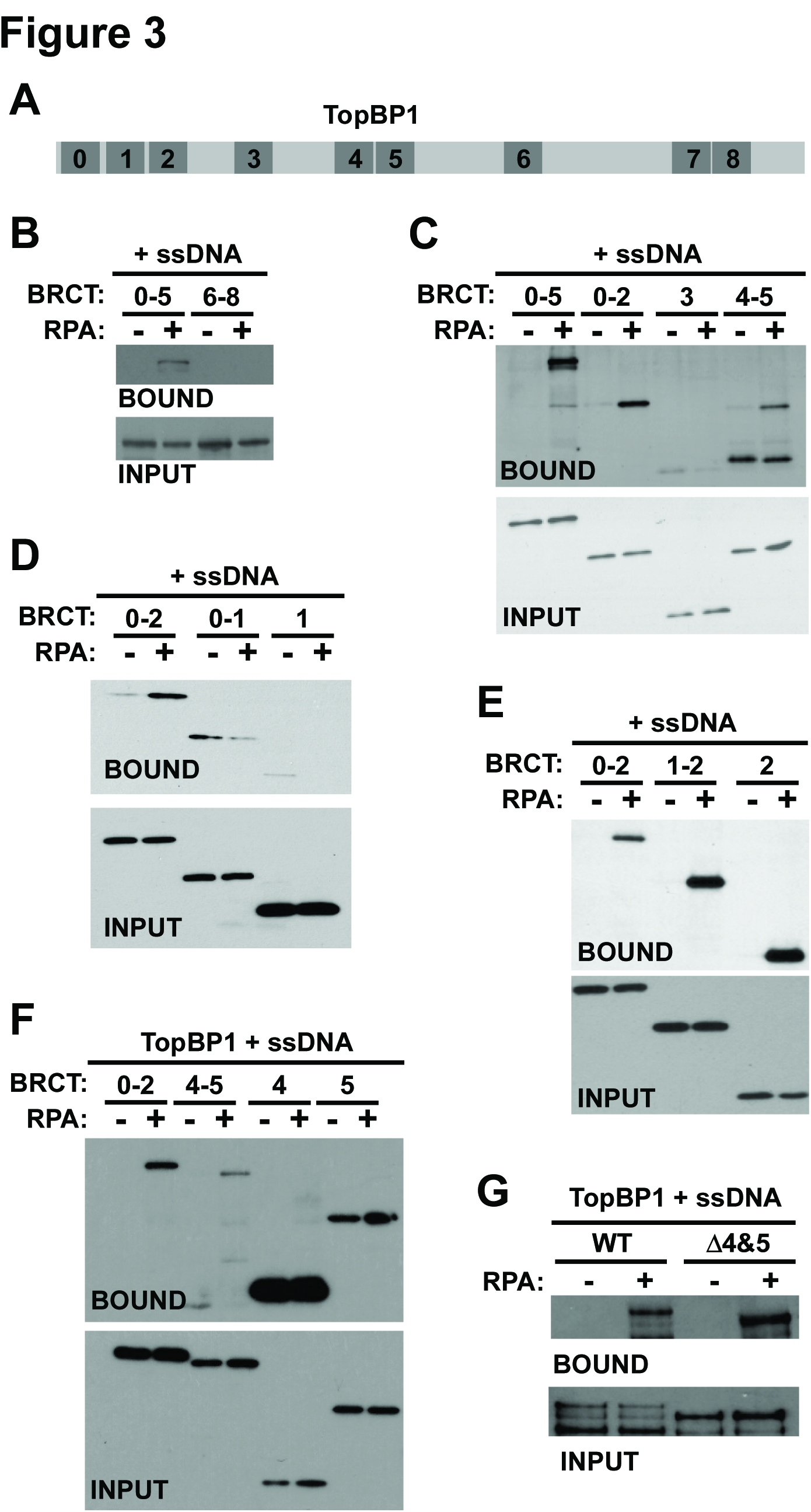
Deletion analysis of TopBP1 binding to RPA-ssDNA. (A) Cartoon depicting the relative positions of BRCT domains (numbered) within the TopBP1 protein. (B-G) Myc-tagged TopBP1 derivatives corresponding to the indicated BRCT domain(s) were produced by IVT and then mixed with biotin-linked ssDNA (1.5 kb) and binding buffer. Recombinant RPA was optionally added, as indicated. After incubation, the ssDNAs were isolated using streptavidin magnetic beads, washed, eluted, and bound proteins detected by Western blot. The TopBP1 proteins were detected using Mab 9E10, which recognizes the myc tag. Input refers to 1.6% of the initial binding reaction.

Our finding that TopBP1 BRCT2 is an RPA-ssDNA binding domain was intriguing given previous work from our laboratory showing that a point mutation within BRCT2, W265R, prevents TopBP1 from accumulating at stalled replication forks, but nonetheless supports replication initiation (1). To pursue this connection, we next asked if TopBP1 W265R could bind RPA-ssDNA using the assays described above. IVT wild type and W265R TopBP1 proteins were mixed with egg extract and either no DNA, dsDNA, or ssDNA, and binding was assessed as in Fig. 1A. As expected, wild type TopBP1 bound to ssDNA, however the W265R mutant did not show detectable binding (Fig. 4A). We note that in this experiment some binding of wild type TopBP1 to dsDNA was observed, which may be due to conversion of some of the dsDNA to ssDNA upon incubation in egg extract. TopBP1 binding to dsDNA was not reproducibly observed (see Fig. 1A). When IVT TopBP1 proteins were mixed with ssDNA and purified RPA, we again observed that wild type, but not W265R TopBP1, could bind ssDNA efficiently, in an RPA-dependent manner (Fig. 4B). We next asked if the W265R mutation in the context of BRCT2 alone would impact binding to RPA-ssDNA. For this experiment, purified recombinant forms of GST-tagged BRCT2, either wild type or with the W265R mutation, were prepared and used for binding assays. As shown in Fig. 4C, GST-BRCT2 bound well to ssDNA in an RPA-dependent manner, while GST-BRCT2 W265R binding is reduced in comparison. This experiment makes two important points. First, because this was a completely purified system, we see that BRCT2 can bind directly to RPA-ssDNA, and furthermore, because all proteins were purified from *E. coli*, no post-translational modifications on either BRCT2 or RPA are required for binding. Second, the data show that W265 is a critical determinant for binding to RPA-ssDNA, both in the context of the full-length protein (Fig. 4A-B) and for BRCT2 alone (Fig. 4C). We next asked if the ssDNA size restrictions observed previously for binding of full-length TopBP1 (Fig. 1B) also applied to the isolated BRCT2 domain, and found this was indeed the case. IVT-produced BRCT2 bound to RPA-ssDNA of 150 nt, but did not bind when the ssDNA was smaller than 150 nt (Fig. 4D).

**Figure 4.**
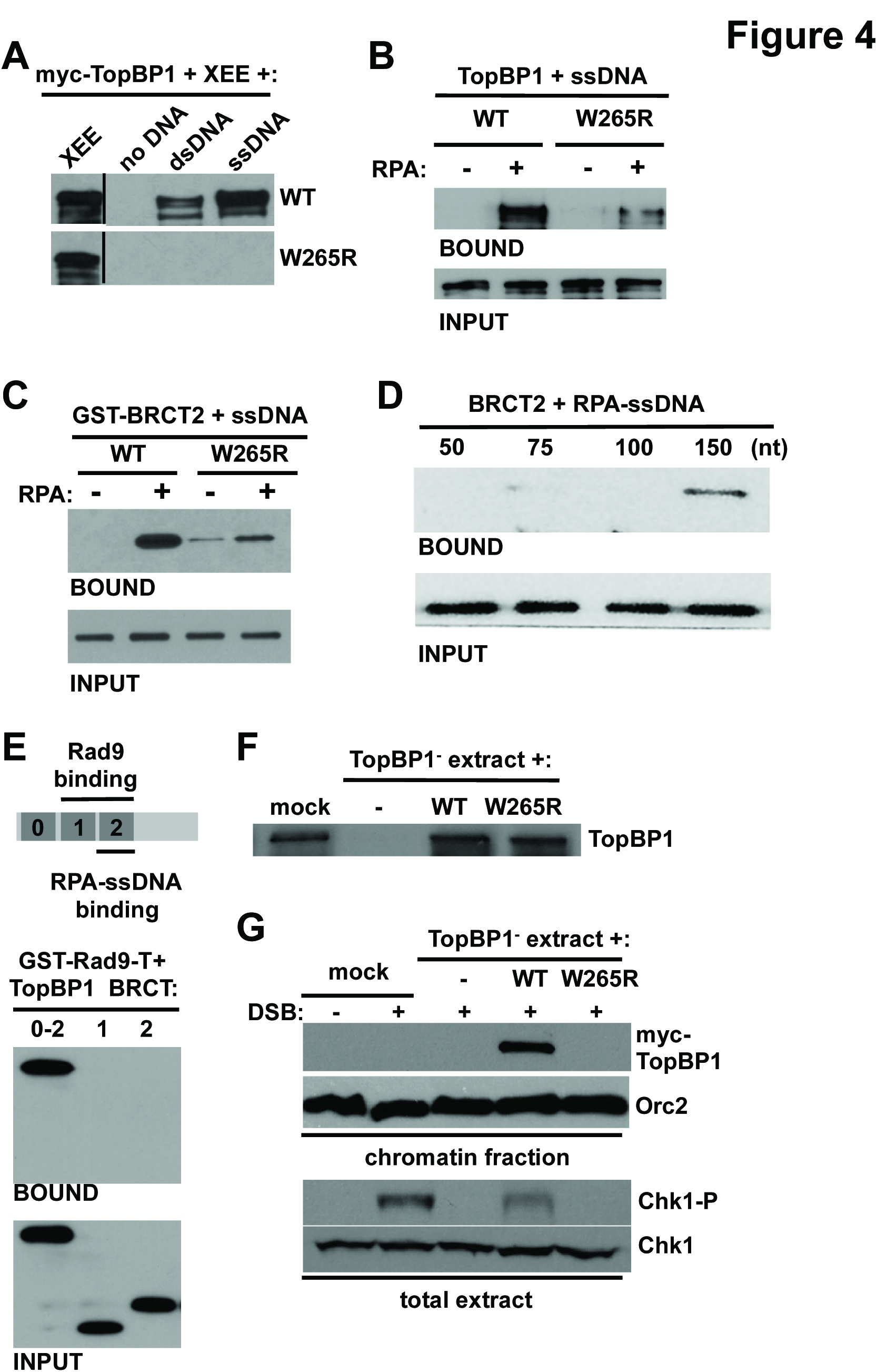
Functional characterization of the TopBP1 BRCT2 interaction with RPA-ssDNA. (A) Myc-tagged full-length TopBP1 proteins, either wild type or the W265R point mutant, were produced by IVT and then mixed with egg extract and the indicated DNAs. Samples were then processed exactly as in Fig. 1A. Proteins were detected with Mab 9E10 against the myc tag. (B) Myc-tagged full-length TopBP1 proteins, either wild type or the W265R point mutant, were produced by IVT and then utilized for the ssDNA binding assay, exactly as described in Figs. 3B-F. (C) Purified, recombinant GST-BRCT2 fusion proteins, either wild type or the W265R mutant, were mixed with biotin-linked ssDNA (1.5 kb) and binding buffer. Recombinant RPA was optionally added, as indicated. After incubation, the ssDNAs were isolated using streptavidin magnetic beads, washed, eluted, and bound proteins detected by Western blot using anti-GST antibodies. Input refers to 1.6% of the initial binding reaction. (D) Myc-tagged TopBP1 BRCT2 fragment was produced by IVT and then mixed with recombinant RPA and biotin-linked ssDNAs of the indicated size. The samples were then processed as in Figs. 2 B-F. Input refers to 1.6% of the initial binding reaction. (E) Cartoon depicting the N-terminal region of TopBP1 and binding determinants for Rad9 and RPA-ssDNA. Myc-tagged TopBP1 fragments, corresponding to the indicated BRCT domains, were produced by IVT and then mixed with purified *Xenopus* Rad9 tail domain fused to GST (GST-Rad9-T). Recombinant casein kinase 2 was also included in the reaction. After incubation, proteins bound to GST-Rad9-T were recovered on glutathione-sepharose beads, washed, eluted, and analyzed by Western blotting. Input refers to 3.3% of the initial reaction volume. (F) Egg extract was either mock depleted (“mock”), or immunodepleted of TopBP1 using TopBP1 antibodies (TopBP1-). TopBP1-depleted extract was then supplemented with either blank IVT (-), or IVTs expressing the indicated myc-tagged TopBP1 protein. The samples were then blotted with antibodies against TopBP1. (G) The egg extracts from (F) were supplemented with sperm chromatin and, optionally, EcoRI to produce DSBs (“+ DSB”). After incubation, the sperm chromatin was isolated as described (1) and the samples blotted with either Mab 9E10, to detect myc-tagged TopBP1, or antibodies against Orc2, as a loading control (panel “chromatin fraction”). In addition, samples of the total extract were blotted for S345-phosphorylated Chk1 (Chk1-P) or Chk1 as a loading control (panel “total extract”).

Previous work has shown that the tandem BRCT1-2 repeats of TopBP1 bind to the phosphorylated Rad9 tail domain (35,37), and we have shown here that BRCT2 alone binds to RPA-ssDNA (summarized in Fig. 4E). The amino-terminal region of the TopBP1 protein thus appears to interact with multiple binding partners that are important for checkpoint signaling. Although previous work had narrowed down the Rad9 binding region of TopBP1 to BRCT1-2 (37), to our knowledge the ability of either isolated BRCT1 or BRCT2 to bind the Rad9 tail domain had not yet been tested. This was of interest, as we wanted to know if the binding determinants within the BRCT1-2 region for Rad9 relative to RPA-ssDNA were identical, or distinct despite being located in the same region of the protein. To examine this, we performed a pull-down assay with a previously described GST-Rad9 tail domain fusion protein (42) and looked for interaction between the Rad9 tail domain and TopBP1 BRCT0-2, BRCT1, or BRCT2. As shown in Fig. 4E, the BRCT0-2 fragment could bind well to the Rad9 tail domain, as expected, however neither BRCT1 or BRCT2 alone could bind. It thus appears that the tandem BRCT1-2 repeats bind Rad9, whereas BRCT2 alone is sufficient to bind RPA-ssDNA. We conclude that distinct determinants control binding of TopBP1 to Rad9 and RPA-ssDNA.

### TopBP1 BRCT2 W265R fails to accumulate on DNA DSB-containing chromatin and fails to activate ATR during a DSB response

Previous work has shown that TopBP1 W265R is not recruited efficiently to stalled replication forks (1), and data shown here demonstrate that this mutant also fails to bind efficiently to RPA-ssDNA (Figs. 4A-C). Taken together, the data strongly suggest that a BRCT2-mediated direct interaction with RPA-ssDNA allows TopBP1 to accumulate on the RPA-ssDNA present at stalled forks. To see if this a general mechanism for how TopBP1 is recruited to sites of DNA damage, we next assessed the ability of TopBP1 W265R to accumulate on chromatin during a DSB response, given that DSB-containing chromatin also contains RPA-ssDNA. Egg extracts were prepared, immunodepleted of endogenous TopBP1, and optionally supplemented with either blank IVT, or IVTs expressing wild type or W265R TopBP1 (Fig. 4F). These extracts were then mixed with sperm chromatin and the EcoRI restriction enzyme, which activates ATR by inducing DSBs (43). After incubation, the chromatin was isolated and probed for myc-tagged TopBP1 proteins and, as a loading control, the Orc2 protein. In addition, samples of the total extract were taken to blot for phosphorylated Chk1 (Chk1-P), as a readout for ATR activity. As shown in Fig. 4G, wild type TopBP1 was bound to the DSB-containing chromatin, however TopBP1 W265R did not bind to DSB-containing chromatin. Furthermore, while wild type TopBP1 could support Chk1 phosphorylation during the DSB response, TopBP1 W265R could not. These data are consistent with our previous work that revealed defects for TopBP1 W265R in associating with stalled forks, and suggest that direct binding to RPA-ssDNA is a general mechanism that allows TopBP1 to be recruited to sites of damage to promote ATR activation.

## DISCUSSION

Data shown here reveal new insights into how DSBs and stalled replication forks are sensed to activate the ATR pathway. In Fig. 5 we present a new model for ATR activation at stalled forks. This model proposes that an early event in ATR activation is the direct interaction between TopBP1 and RPA-ssDNA at the stalled fork, via TopBP1’s BRCT2 domain. TopBP1 binding then allows recruitment of Rad17 and 911 (1), perhaps through an interaction that changes the conformation of Rad17, thereby allowing it to more productively interact with the 5′-DNA junction (as depicted in the figure), or through clearance of other factors that might preclude interaction between Rad17 and the 5′-DNA junction. 911 loading then ensues, and we propose that this event stabilizes TopBP1 at the stalled fork, possibly because TopBP1 moves from its binding site on RPA-ssDNA to a higher-affinity binding site on the Rad9 tail domain (as depicted in the figure), or because additional molecules of TopBP1 join the complex via binding to Rad9. Stabilization of TopBP1 at the stalled fork after clamp loading is consistent with previous work showing that TopBP1 recruitment to stalled forks is reduced (but not eliminated) upon Rad17 depletion (33,36). Finally, once TopBP1 is bound to Rad9, it can interact with the independently recruited ATR-ATRIP complex, and ATR is activated. We note that this model does not include the MRN complex, which others have shown to be involved in ATR activation (31,33,42), as it is still unclear if MRN plays a direct role in activation, or acts indirectly through the production of RPA-coated ssDNA. In summary, our new data, combined with previous results, show that stalled forks are independently recognized by two sensors, ATRIP and TopBP1, and that a common mechanism is employed: direct interaction between these sensors and RPA-ssDNA. These findings underscore the importance of RPA-ssDNA as a platform for checkpoint complex assembly and activation.

**Figure 5.**
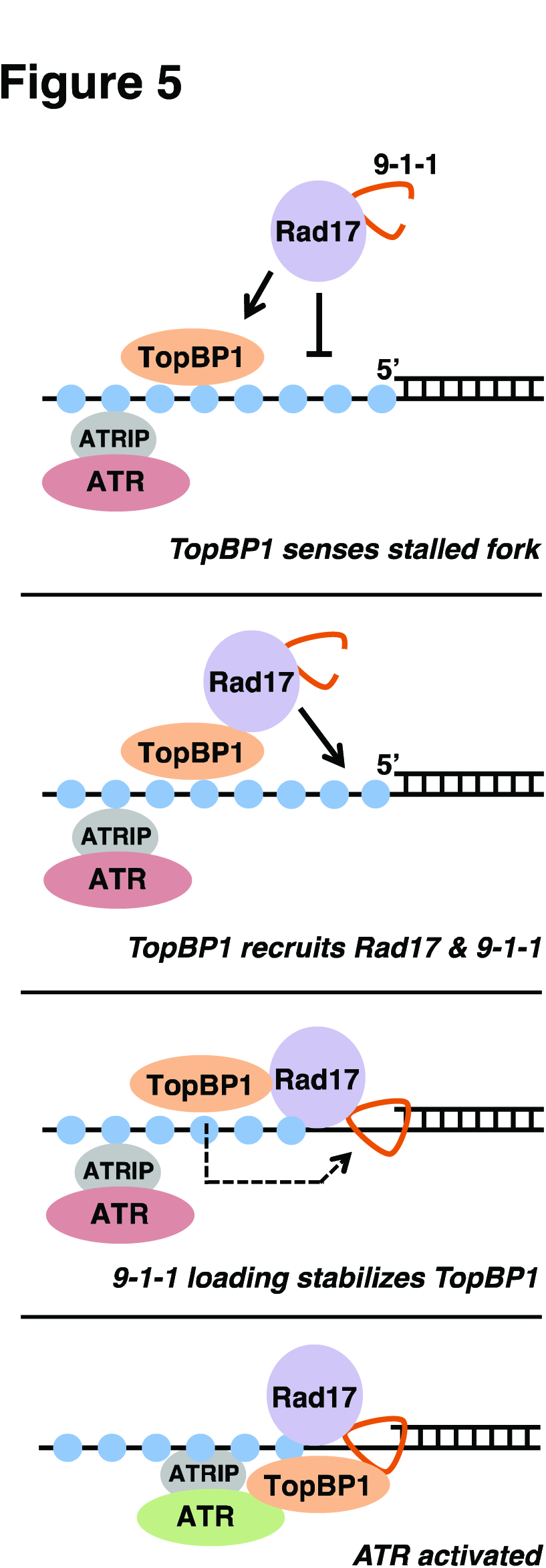
ATR activation during a checkpoint response. A new model for ATR activation, please see main text for details.

## Acknowledgements

We thank Oscar Aparicio, Irene Chiolo, and members of our laboratory for comments on the manuscript. This work was funded by NIH R01GM099825 to W.M.M.

## AUTHOR CONTRIBUTIONS

J.A., S.Y., and W.M.M. performed the experiments. W.M.M. proposed and supervised the project, contributed to research design, and analyzed the results. J.A. designed the study, performed most experiments, and analyzed the results. The paper was co-written by J.A. and W.M.M.

